# Cryptochrome Loss Drives COPD-like Lung Pathology through Disrupted Alveolar Epithelial Proliferation and Immune Homeostasis

**DOI:** 10.64898/2026.05.19.726266

**Authors:** Tingting W. Mills, Chorong Han, Ji Ye Lim, Nobuya Koike, Sun Young Kim, Jaebok Wi, Hui Liu, Yu Wang, Kazuhiro Yagita, Joseph L. Alcorn, Zheng Chen, Seung-Hee Yoo

## Abstract

Chronic obstructive pulmonary disease (COPD) is a progressive lung disease characterized by alveolar destruction, impaired epithelial regeneration and chronic inflammation. While circadian disruption has been linked to COPD pathogenesis, the cellular mechanisms remain unclear. Here, we identify the core clock components Cryptochrome 1 and 2 (*Cry1/2*) as essential regulators of alveolar epithelial cell proliferation and pulmonary immune homeostasis. *Cry1/2* double knockout (dKO) mice exhibit spontaneous emphysema-like pathology, including airspace enlargement, increased lung compliance, and inflammatory cell infiltration without environmental insults. Mechanistically, *Cry1/2*-deficient alveolar epithelial cells display reduced proliferative capacity, and transcriptomic profiling revealed a pronounced shift toward a more proximal airway-like phenotype. Notably, these cells share a gene signature with human COPD lungs in pathways involving immune regulation. Furthermore, *Cry1/2*-deficient macrophages show elevated responsiveness to LPS. Bone marrow (BM) transplantation revealed that mice receiving *Cry1/2* dKO BM suffer from enhanced lung inflammation without airspace enlargement, supporting a critical role of epithelial *Cry1/2* in maintaining alveolar integrity. Importantly, treatment with Nobiletin, a circadian rhythm-modulating compound, mitigates NF-κB activation and ameliorates lung inflammation and structural damage in *Cry1/2* dKO mice. These findings establish CRY1/2 as critical circadian regulators of epithelial cell proliferation and immune homeostasis, and highlight the therapeutic potential of targeting circadian pathways in COPD.

## Introduction

Chronic obstructive pulmonary disease (COPD) is a progressive and life-threatening respiratory disorder and remains a major public health concern. It is the third leading cause of death worldwide, with a global prevalence of approximately 11.7% (95% CI: 8.4-15%)^1, 2, 3, 4^. COPD is characterized by persistent airway inflammation, epithelial cell damage, and excessive mucus production, ultimately leading to airflow limitation, reduced lung function, and respiratory distress. While significant advancements have been made in identifying genetic and environmental risk factors, such as α1-antitrypsin deficiency and smoking, and existing treatments can alleviate symptoms and slow disease progression ^5, 6, 7, 8, 9, 10, 11, 12, 13^, there is currently no cure for COPD^14^. Therefore, an in-depth understanding of the molecular mechanisms driving COPD pathogenesis is essential for developing effective therapeutic strategies.

The mammalian circadian clock is an intrinsic timekeeping system composed of cell-autonomous molecular oscillators present in nearly all cells of the body. At its core, the circadian machinery operates through transcriptional-translational feedback loops involving both positive regulators including Circadian Locomotor Output Cycles Kaput (CLOCK), Brain and Muscle ARNT-Like 1 (BMAL1), and Retinoic Acid Receptor-Related Orphan Receptors (RORs), and negative regulators including Period (PER1, PER2, PER3), CRYPTOCHROMES (CRY1, CRY2), and REV-ERBs (15–17). The heterodimeric CLOCK/BMAL1 complex initiates the transcription of target genes, including PER1/2/3 and CRY1/2 (15, 16). Once translated, PER and CRY proteins form a heterodimer that translocates back into the nucleus and inhibits CLOCK/BMAL1 activity and thereby their own transcription (17). This core feedback loop is further stabilized by an interlocked secondary feedback loop where the competing nuclear receptors REV-ERBs and RORs regulate the timing and amplitude of BMAL1 expression ^15^. Through these tightly controlled molecular interactions, the circadian clock drives rhythmic gene expression in alignment with 24-hour environmental cycles to maintain physiological homeostasis ^16, 17, 18, 19, 20^.

Growing evidence indicates that circadian clocks regulate essential functions in the lung, such as forced expiratory volume in 1 second (FEV₁), the FEV₁/forced vital capacity (FVC) ratio, airway resistance, immune cell composition, and inflammatory responses ^21, 22^. At the cellular level, lung epithelial cells (ECs) exhibit high expression of core clock genes, suggesting a potential role of the circadian clock in lung homeostasis ^23, 24, 25, 26, 27^. In accordance, genetic deletion of the core clock gene *Bmal1* in bronchial ECs disrupts the circadian rhythm of neutrophil recruitment ^25^, and the circadian clock in alveolar ECs contributes to the repair process and inflammatory response after cigarette smoke (CS) stimulation ^28^. Furthermore, clinical and epidemiological studies have demonstrated that circadian misalignment, which is common in shift workers and those with sleep disorders, is associated with increased COPD prevalence and severity ^29^. COPD patients often report exacerbated symptoms during early morning and nighttime hours, which coincide with the circadian nadir in lung function ^30, 31, 32^. Consistently, deterioration of sleep quality and reduction in melatonin ^33^ are among the frequent features in COPD patients and have been linked to increased exacerbations, emergency healthcare utilization, and mortality ^34, 35, 36^. Finally, animal studies have shown that mice subjected to both acute and chronic exposure to CS, one of the major causes of COPD, have dysregulated oscillation and expression of core clock genes and their targets in the lungs ^37, 38^.

Despite this growing body of literature linking circadian rhythms to pulmonary function and immune regulation, the molecular mechanisms connecting circadian disruption to COPD pathogenesis remain largely unclear. In this study, we aim to identify and characterize key circadian components in regulating lung inflammation and protecting against emphysema in COPD. By investigating how circadian control influences epithelial and immune cells, we seek to uncover novel molecular pathways driving disease progression and evaluate the potential of clock-targeted chronotherapeutic strategies to mitigate COPD severity.

## Materials and methods

### Mouse Generation and Treatment

All mice used in this study were on the C57BL/6J genetic background. Animals were housed under pathogen-free conditions at the University of Texas Health Science Center in Houston, TX (UTHealth Houston). All experimental procedures were approved by the UTHealth Houston Animal Welfare Committee. *Cry1* and *Cry2* double knockout (*Cry1/2* dKO) mice and *Per1* and *Per2* double knockout (*Per1/2* dKO) mice, reported previously ^39, 40^, were used in this study. For the LPS-induced COPD-like model, three-month-old WT or *Cry1/2* dKO mice were injected with lipopolysaccharide (LPS, 0.2 mg/kg; Millipore-Sigma, L4391) through oropharyngeal aspiration twice per week for four weeks. On day 30, mice were euthanized, and lungs were harvested for histological and molecular analysis. For the Nobiletin study, mice were fed with standard chow alone or standard chow supplemented with 0.1% NOB (10 mg/kg b.w.) beginning at weaning (postnatal day 25) and continued for 20 weeks.

### Cell culture

Human lung epithelial cell line A549 and mouse primary lung epithelial cell line MLE12 were procured from ATCC (Manassas, VA). Cells were maintained in a 37°C incubator supplemented with 5% CO₂. A549 cells were cultured in Dulbecco’s Modified Eagle Medium (DMEM) supplemented with 10% fetal bovine serum (FBS) and 1% antibiotics. MLE12 cells were maintained in 1:1 Ham’s F12/DMEM medium supplemented with 2% FBS, 0.005 mg/ml insulin, 0.01 mg/ml transferrin, 30 nM sodium selenite, 10 nM hydrocortisone, 10 nM β-estradiol, 2 mM L-glutamine, and 1% antibiotics. For siRNA-mediated knockdown, cells were cultured in antibiotic-free medium and transfected with 50 pmol/ml of *CRY1, CRY2, CRY1/2*, or control siRNA (Sigma) using Lipofectamine® RNAiMAX (Life Technologies) on days 0 and 1. Cells were collected on day 3 for assays. For the WST-1 (Water-Soluble Tetrazolium salt-1) assay, cells were cultured in 96-well plate, transfected with siRNA as described above, and incubated with WST-1 reagent (Roche) for 4 hours. Absorbance was measured at 450 nm using a microplate reader to assess cell viability.

### Alveolar epithelial cell (EC) isolation

Primary human lung ECs were isolated as previously described using dispase digestion ^41, 42^. Mice were anesthetized with 2.5% Avertin, and lungs were perfused with cold PBS. Dispase (5 U/ml, StemCell Technologies) was injected through the trachea in 0.5–1 ml aliquot over 45 minutes (total ∼2 ml/mouse). After digestion, lung lobes were separated, minced into ∼1 mm³ pieces, and incubated in epithelial growth medium (DMEM with 10% FBS and 1% penicillin-streptomycin antibiotics) with 50 U/ml DNase I (Millipore Sigma) for 15 minutes. After filtering through a 40 µm strainer, immune cells were depleted using anti-CD16/32 and anti-CD45 antibodies, followed by magnetic bead separation (BD Biosciences). Isolated epithelial cells were cultured in epithelial growth medium at 37°C in a humidified incubator with 5% CO₂. After 2–3 days, non-adherent cells were removed, and the adherent epithelial cells were harvested for RNA, protein, and RNA-seq analysis.

### Lung Function Assay

Pulmonary function was assessed using the flexiVent FX system (SCIREQ) with the FX1 module and Flexiware v8.3 software. Mice were anesthetized via intraperitoneal injection of 2.5% Avertin (0.012 ml/g). A tracheostomy was performed, and lungs were ventilated via an 18-gauge cannula. Lung mechanics were measured using forced oscillation techniques, including compliance (Crs), elastance (Ers), Newtonian resistance (Rn), tissue damping (G), tissue elastance (H) and inspiration capacity (IC). Pressure-volume curves were generated using a ramp maneuver to 30 cmH₂O. Three measurements were taken per mouse, and the average was used for analysis.

### Inflammation and Cellular Differential Assay

BAL fluid was obtained by lavaging the lungs three times with 0.4 ml sterile PBS. Total BAL cell counts were determined using a hemocytometer. Cytospins were prepared by centrifugation (1200 rpm, 5 minutes) and stained with Diff-Quick (Dade Behring) to identify immune cells. Percentages of macrophages, lymphocytes, and neutrophils were calculated and multiplied by the total cell counts to determine absolute numbers. Analyses were performed with blinding to experimental conditions.

### Western Blotting

Lung tissues were pulverized in liquid nitrogen and lysed in RIPA buffer (Boston Bioproducts). Proteins were separated on 4–20% SDS-PAGE gels (Bio-Rad), transferred to PVDF membranes (Millipore Sigma), and blocked with EveryBlot Blocking Buffer (Bio-Rad). Membranes were incubated overnight at 4°C with primary antibodies against CRY1, CRY2 ^43^, β-actin (Sigma), or GAPDH (Life Technologies), C2 (Cell Signaling Technology), C4b (Cell Signaling Technology), and P65 (Cell Signaling Technology) followed by corresponding HRP-conjugated secondary antibodies (Cell Signaling Technology). Signals were visualized with ECL (GE Healthcare) and imaged using a Bio-Rad Chemidoc.

### Real-time PCR and sequencing

Total RNA was extracted using the RNeasy Mini Kit (Qiagen), including on-column DNase treatment. For qPCR, cDNA was synthesized using Gendepot reverse transcription supermix, and reactions were performed on a Bio-Rad CFX384 using gene-specific primers (Supplementary Table 2). Expression levels were normalized to β-actin or 18S rRNA using the ΔΔCt method. RNA sequencing was performed by Novogene using the total RNA, and libraries were prepared with the NEBNext Ultra II RNA Library Prep Kit. The sequencing was performed on an Illumina NovaSeq platform. Differential gene expressions were analyzed using DESeq2, and pathway enrichment was performed using Metascap. The raw RNA-sequencing data have been deposited in the Gene Expression Omnibus (GEO) under accession numbers GSE313652 and GSE313683.

### Bone marrow derived macrophage (BMM) culture

Bone marrow cells were isolated from femurs of euthanized mice by flushing with cold sterile PBS using a 25G needle. The cell suspension was passed through a 70 µm cell strainer to remove debris and centrifuged at 300 × g for 5 minutes. The pellet was resuspended and plated in non-tissue culture-treated Petri dishes in DMEM supplemented with 10% FBS, 1% antibiotics, and 20% L929-conditioned medium, which provides macrophage colony-stimulating factor (M-CSF) to promote macrophage differentiation. Cells were incubated at 37°C with 5% CO₂, and fresh medium was replenished on day 3. On day 7, adherent BMMs were collected and plated for downstream experiments. For inflammatory stimulation, BMMs were treated with LPS (1 μg/mL) for 24 hours. Total RNA was then extracted for real-time qPCR analysis to assess cytokine expression.

### Tissue embedding

Formalin-fixed lungs were washed in PBS and dehydrated through a graded ethanol series (30%, 50%, 70%, 85%, 95%, and 100%), with each concentration applied twice for 30 minutes per step. Dehydrated tissues were then cleared with a 1:1 solution of Histo-Clear (National Diagnostics) and absolute ethanol for 20 minutes, followed by two changes of 100% Histo-Clear for 20 minutes each to remove residual alcohol. The cleared tissues were infiltrated with molten paraffin wax at 60°C in a vacuum oven for 30 minutes, with two paraffin changes to ensure complete infiltration. Tissues were then oriented and embedded in fresh paraffin in embedding molds and allowed to solidify at room temperature. Paraffin blocks were sectioned at a thickness of 4 µm using a Leica RM2235 microtome and mounted onto positively charged glass slides for subsequent histological and immunohistochemical analysis.

### Hematoxylin and Eosin (H&E) Staining and Mean Linear Intercept Analysis

Sections were deparaffinized in Histo-Clear and rehydrated through a graded ethanol series. Tissues were stained with hematoxylin (Epredia™) for 5 minutes, rinsed, and blued in 0.1% ammonia water. Slides were counterstained with eosin (Epredia™) for 30 seconds, dehydrated, cleared, and coverslipped. For MLI analysis, images were captured from the H&E-stained section under 10× magnification using a brightfield microscope. For each mouse, 8∼10 randomly selected, non-overlapping fields were analyzed. A grid of parallel lines was superimposed on each image using ImageJ software, and the number of alveolar wall intercepts per unit length of line was counted. MLI was calculated by dividing the total length of test lines by the number of intercepts, providing an estimate of the average distance between alveolar walls. All measurements were performed in a blinded manner to ensure objectivity.

### Statistics

Data were first assessed for normality using the Shapiro-Wilk Test. For normally distributed data, two-group comparisons were made using a two-tailed Student’s t-test. For comparisons involving more than two groups, ANOVA was followed by post hoc multiple comparisons. Non-parametric tests were used where normality assumptions were not met. A p-value < 0.05 was considered statistically significant. Results are presented as mean ± standard error of the mean (SEM).

## Results

### Cry1/2 double knockout (dko) mice exhibited airspace enlargement in the lung

To determine the impact of clock disruption on lung function, we first examined lung phenotypes in two mouse models harboring deletion of essential clock components, namely *Per1/2* double knockout (dKO) and *Cry1/2* dKO mice. Whereas no abnormal lung histology and no difference in lung function were observed in *Per1/2* dKO mice (Supplementary Fig. 1A-1C), *Cry1/2* dKO mice exhibited airspace enlargement in both male and female mice compared to wildtype (WT) mice (Fig. 1A). This observation is particularly interesting, as previous reports have shown emphysema in other circadian mutants only following environmental challenges such as CS or LPS exposure ^44, 45^. In contrast, *Cry1/2* dKO mice spontaneously develop emphysema-like pathology without such triggers, suggesting a unique and essential role for CRYs in maintaining lung structure. Immunostaining confirmed that both CRY1 and CRY2 are expressed in alveolar epithelial cells (Fig. 1B). To quantify alveolar enlargement, the mean linear intercept (MLI) was measured and found to be significantly elevated in *Cry1/2* dKO mice compared to WT mice (25.8±1.7 μm vs 18.5± 1.4 μm) (Fig. 1C). A mild increase in MLI and lung compliance, as well as decreased lung elasticity, was also observed in *Cry1* or *Cry2* single knockout mice, suggesting partially redundant roles for CRY1 and CRY2 in protecting against emphysema (supplementary Fig. 1D-1F). Inflammatory profiling revealed elevated neutrophils in the bronchoalveolar lavage (BAL) fluid of *Cry1/2* dKO mice (Fig. 1D), indicating augmented airway inflammation—another hallmark of COPD. Consistent with these findings, the lung functional assay with flexiVent showed increased compliance (Crs), decreased elasticity (Ers), and increased inspiration capacity (IC) in *Cry1/2* dKO mice (Fig. 1E), indicating airspace enlargement (emphysema). Because α1-antitrypsin (A1AT) deficiency is a hereditary cause of COPD, particularly emphysema due to the lack of inhibition of neutrophil elastase, we examined the transcript levels of genes coding A1AT in the livers, the major source of the circulating A1AT, from WT and *Cry1/2* dKO mice. Notably, the highly expressed A1AT paralogs (*Serpina1b*, *Serpina1d*, and *Serpina1e*) did not differ between WT and *Cry1/2* dKO mice in either gender (Supplementary Fig. 2). In contrast, paralogs with lower expression showed variable changes as a function of CRY and gender. For example, *Serpina1a* was decreased in female *Cry1/2* dKO mice, and *Serpina1c* was downregulated and *Serpina1f* was upregulated in both male and female *Cry1/2* dKO mice (Supplementary Fig. 2). Therefore, it is unlikely that the emphysema phenotype observed in *Cry1/2* dKO mice is due to α1-antitrypsin deficiency, suggesting a mechanism involving other paralogs. Taken together, our data demonstrate that *Cry1/2* depletion in mice is sufficient to drive both inflammation and the emphysema-like phenotype in the lung.

**Figure 1.**
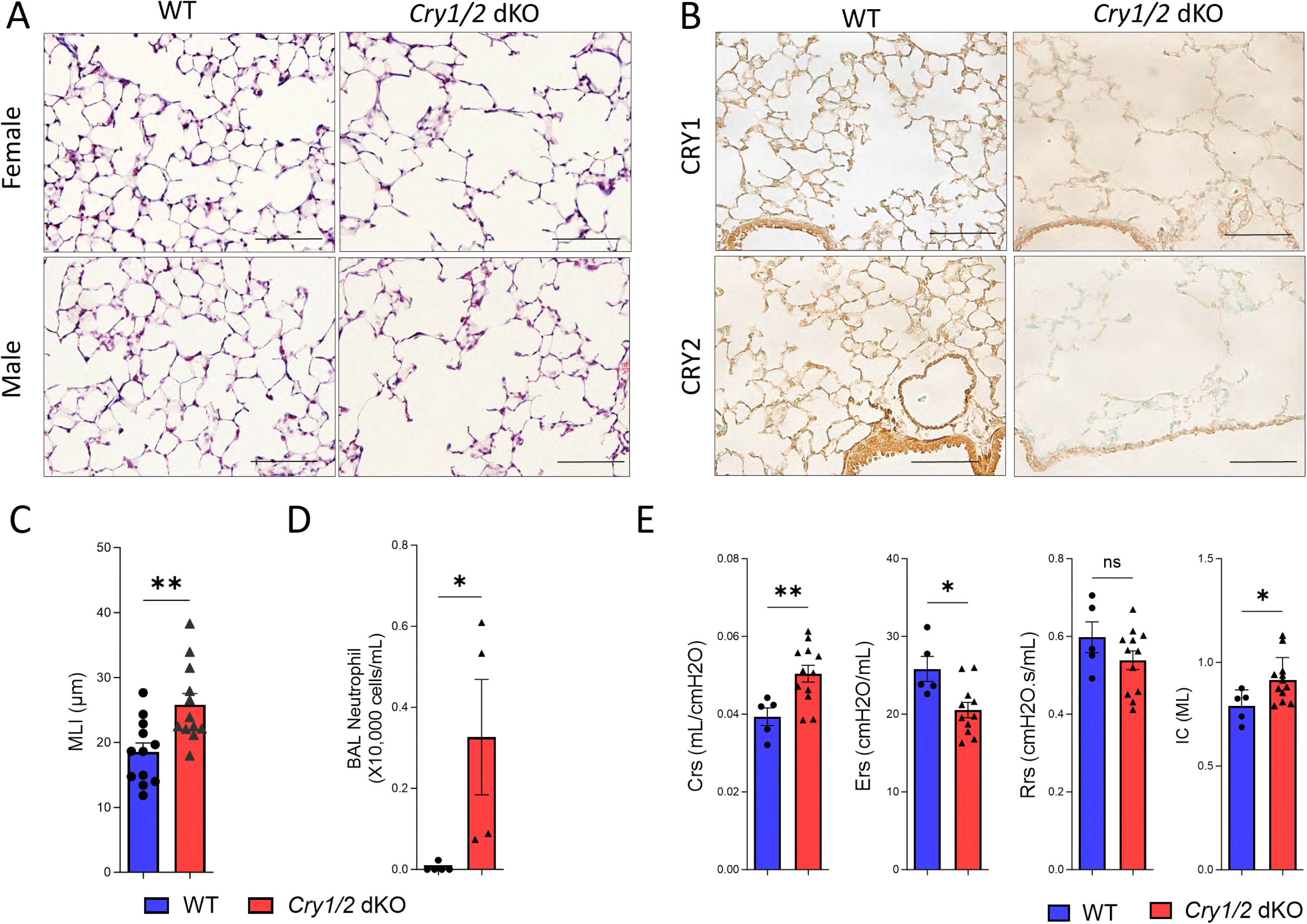
CRY1 and CRY2 double knockout (*Cry1/2* dKO) mice developed spontaneous emphysema. The lungs were collected from 10-12-week old (**A-F**), sex-matched wildtype (WT) and *Cry1/2* dKO mice for analysis. A. Representative images of hematoxylin and eosin (H&E) stained WT and *Cry1/2* dKO lung sections collected from 10-12-week-old male or female mice. **B.** Representative CRY1 and CRY2 antibody (brown) stained lung sections of 10-12-week-old WT and *Cry1/2* dKO mice. Scale bar=100 μm. **C.** Quantification of airspace enlargement by mean linear intercept (MLI) analysis in 10–12-week-old mice (n=12). **D.** Neutrophil counts in bronchoalveolar lavage (BAL) fluid. **E**. Lung function measurements obtained using the flexiVent system, including compliance, elasticity, and inspiratory capacity. Data are presented as mean ± SEM. P-value was determined by an unpaired two-tailed Student’s t-test. * p<0.05, ** p<0.01.

### CRY2 is downregulated in a mouse model of chronic lipopolysaccharides (LPS)-induced lung injury

To examine whether CRYs are dysregulated in COPD, we employed a chronic LPS mouse lung injury model ^46, 47^. In this model, mice were injected with LPS at a low dose (0.2 mg/kg) twice a week for 4 weeks, and the lungs were analyzed on day 30 (Fig. 2A). The LPS injected lungs displayed increased inflammatory cell infiltration (Fig. 2B) and enhanced expression of cytokines including monocyte chemoattractant protein-1 (Ccl2), chemokine (C-X-C motif) ligand 9, 10 and 11 (*Cxcl9*, *Cxcl10* and *Cxcl11*), transforming growth factor-β (*Tgfb*) and Tumor necrosis factor alpha (*Tnfa*) (Fig. 2C). Interestingly, CRY2 protein levels were significantly downregulated in the lungs of LPS-treated mice, while CRY1 protein levels remained unchanged (Fig. 2D). These findings suggest that CRY2 is dysregulated in an animal model of emphysema and its downregulation may directly impact disease pathogenesis.

**Figure 2.**
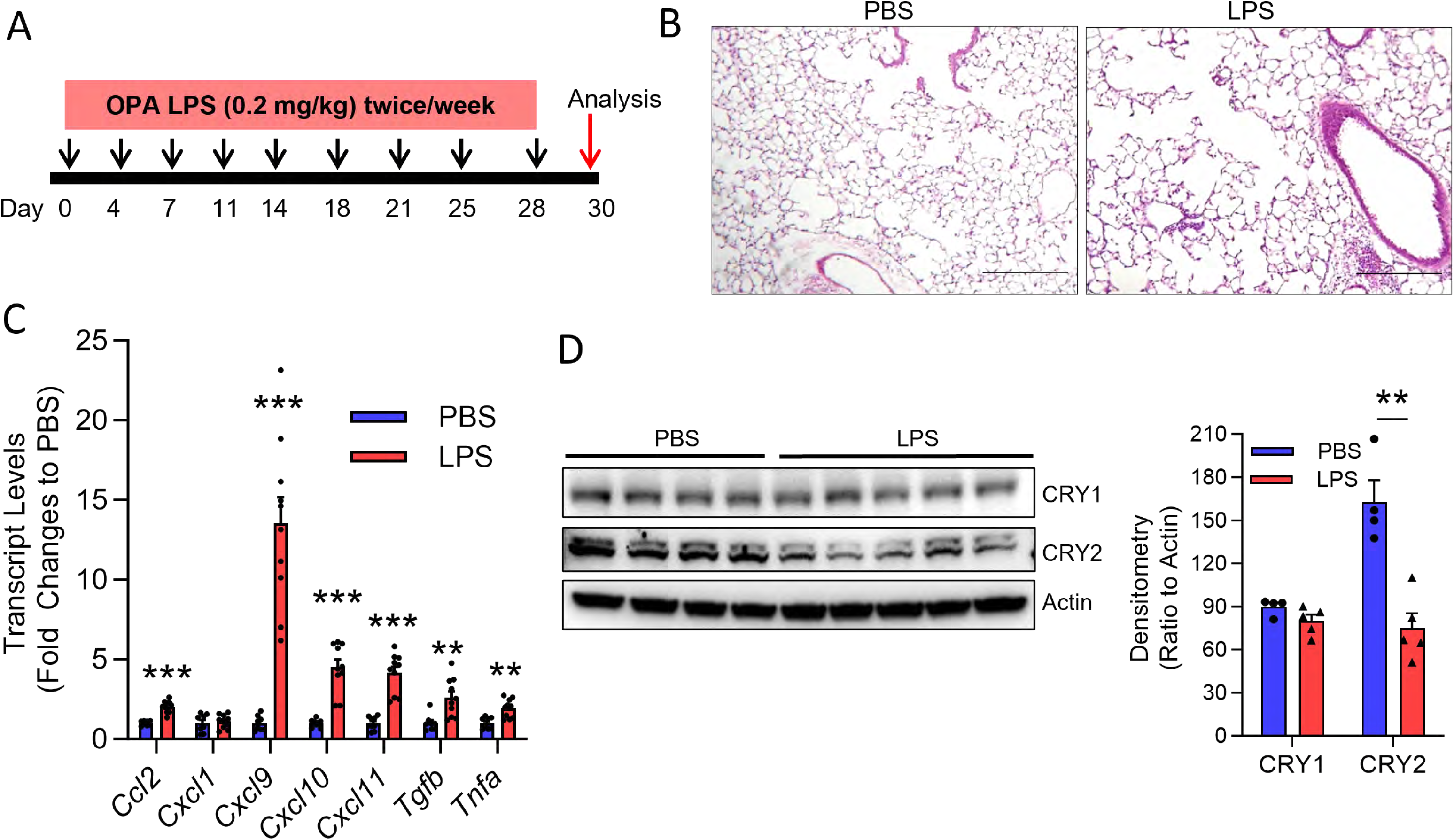
CRY1 and CRY2 expressions in chronic LPS-induced lung injury. 10-12-week-old WT mice were injected with 0.2 mg/kg LPS through oropharyngeal aspiration (OPA) twice a week for 4 weeks. The lungs and BALs were collected on day 28 for analysis. **A.** Schematic diagram of the experimental schedule. **B.** Representative H&E images of WT mice treated with PBS or LPS. Scale bar=200 μm. **C.** RT-qPCR showing the transcript levels of cytokines in the lungs of mice treated with PBS or LPS. **D.** Right panel: The protein levels of CRY1 and CRY2 were determined by western blot. Left panel: Densitometry analysis of the western blot image. Data are presented as mean ± SEM. **p<0.01, ***p<0.001, LPS vs PBS based on Bonferroni adjusted t-test.

### *Cry1/2* deletion promotes LPS-induced emphysema-like pathology and inflammation

Next, to determine whether CRYs are protective in LPS-induced emphysema pathology, we subjected WT or *Cry1/2* dKO to chronic LPS (Fig. 2A) to induce COPD-like phenotypes. Compared to WT mice, *Cry1/2* dKO mice showed a marked enrichment of BAL lymphocytes in response to LPS (Fig. 3A). The BAL neutrophil numbers were highly elevated in both groups compared to the baseline levels in Fig. 1D, but did not differ between WT and *Cry1/2* dKO mice, possibly because chronic LPS induces a ceiling neutrophil response that overrides genotype-specific differences. Lung function analysis revealed that compliance and inspiration capacity were significantly enhanced, while the elasticity was impaired in *Cry1/2* dKO mice compared to WT mice (Fig. 3B), suggesting a more severe emphysema phenotype in *Cry1/2* depleted mice. Consistently, H&E-stained slides showed greater inflammation in *Cry1/2* dKO lungs compared to WT (Fig. 3C). These results suggest that the deletion of *Cry1/2* aggravates LPS-induced COPD phenotypes in mice by promoting emphysema pathology and inflammation.

**Figure 3.**
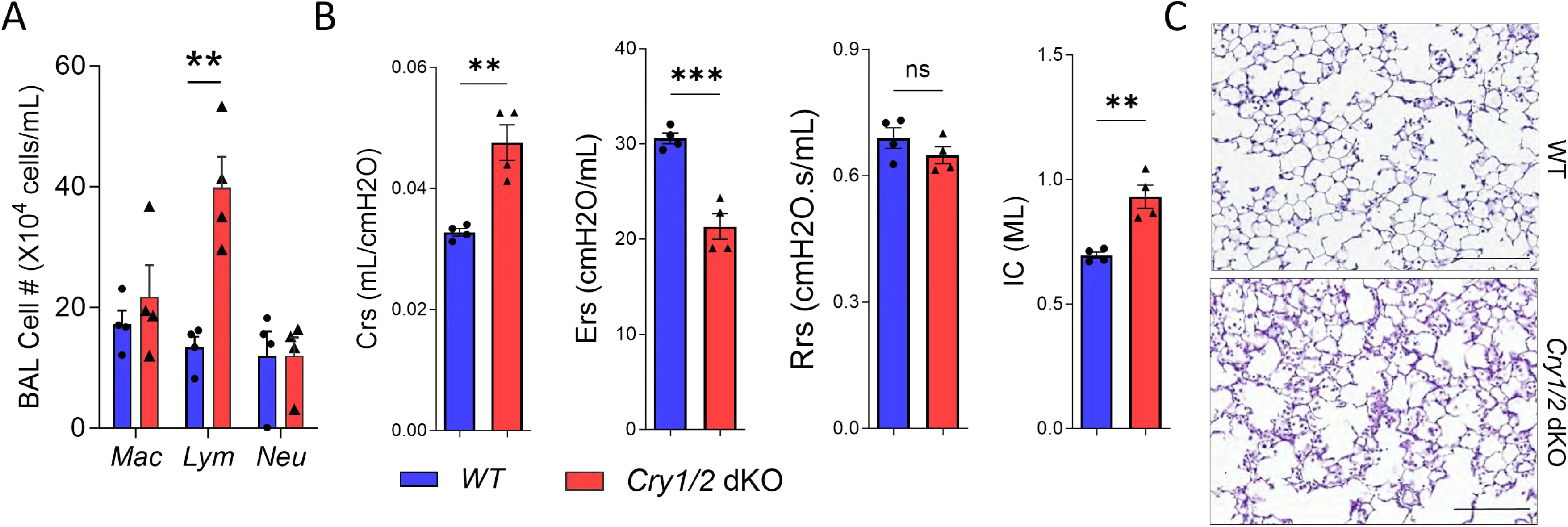
*Cry1/2* depletion promotes LPS-induced lung inflammation. Age and sex-matched WT and *Cry1/2* dKO mice were treated with LPS (0.2 mg/kg) via OPA twice a week for 4 weeks and analyzed on day 28. **A.** The immune cell counts in the BAL fluid. **B.** Lung function measurements obtained using the flexiVent system. **C.** Representative H&E images of WT or *Cry1/2* dKO mice treated with LPS. Scale bar=200 μm. Data are presented as mean ± SEM. ** p<0.01, *** p<0.001 *Cry1/2* dKO vs WT based on Bonferroni adjusted t-test.

### *Cry1/2* depletion in lung epithelial cells exacerbates the inflammatory response and disrupts cell proliferation

The alveolar epithelial cells (ECs) play an important role in maintaining lung hemostasis through surfactant production, barrier protection, fluid clearance, initiation of immune responses, and injury repair. Conversely, alveolar EC death/senescence or dysfunction significantly contributes to COPD and emphysema pathogenesis^48, 49, 50^. To determine whether *Cry1/2* deletion facilitates the emphysema-like phenotype by altering epithelial cell function, we performed *in vitro* studies using the MLE12 murine alveolar epithelial cell line. Cells were transfected with siRNAs targeting *Cry1*, *Cry2*, or both, and subsequently assessed for cytokine production (Fig. 4A). Knockdown of *Cry1/2* led to increased expression of proinflammatory cytokines, including C-C motif chemokine ligand 2 (*Ccl2*), C-X-C motif chemokine ligand 1 (*Cxcl1*), Interleukin-8 (*Il8*, *Cxcl15* in mouse), and *Tnfa* (Fig. 4B). Additionally, *Cry1/2* knockdown cells displayed significantly reduced activities in WST-1 assays, suggesting impaired cell proliferation (Fig. 4C). Similar results were observed in A549 human lung epithelial cells, where knockdown of *CRY2*, but not *CRY1,* resulted in diminished cell numbers (Supplementary Fig. 3), possibly due to CRY2 playing a dominant role in regulating proliferation-related pathways in these cells.

**Figure 4.**
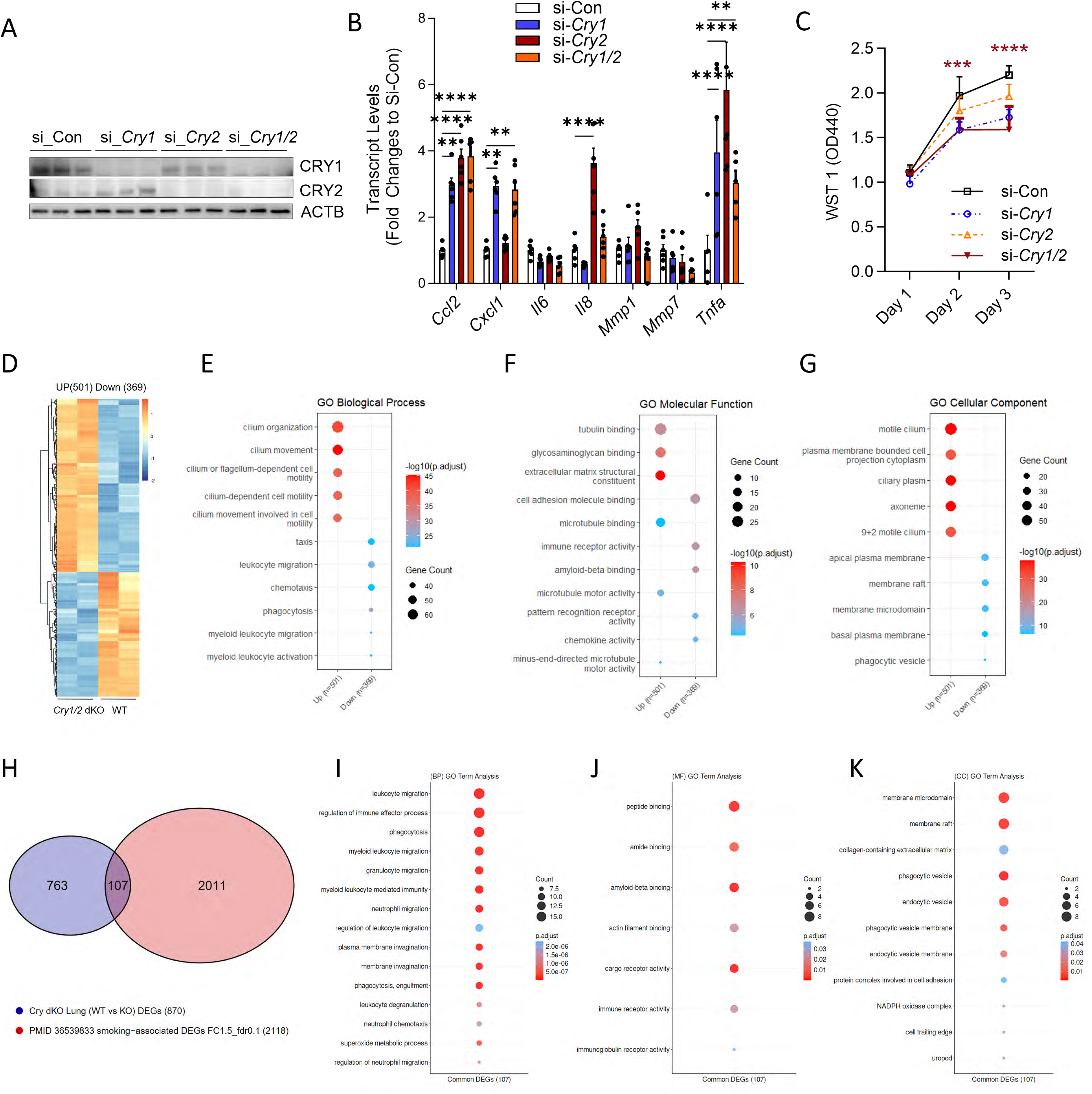
The role of CRY1 and CRY2 in lung epithelial cells. MLE12 cells were transfected with 50 pmole/ml control, *Cry1*, or *Cry2* siRNA on day 0 and day 1. **A.** Western blot showed CRY1/CRY2 protein levels on day 2. **B.** The transcript levels of cytokines, including C-C motif chemokine ligand 2 (*Ccl2*), C-X-C motif chemokine ligand 1 (*Cxcl1*), Interleukin-6 (*Il6*), *Il8/Cxcl5*, matrix metalloproteinase 1 and 7 (*Mmp1* and *Mmp7*), and *Tnfa* were determined with RT-qPCR. **C.** Cell numbers were measured with the WST1 assay 1-, 2-, or 3-days post siRNA transfection. n=6. **D-E.** Primary alveolar ECs were isolated from WT or *Cry1/2* dKO mice for RNA sequencing analysis. D. Heatmap image of up (501) and down (369) regulated genes in *Cry1/2* dKO mice alveolar ECs compared to WT alveolar ECs. Each gene is represented as a horizontal line. E. GO Biological Process (BP) analysis of DEGs that were in *Cry1/2* dKO lung ECs. **F.** GO Molecular Function (MF) analysis of DEGs. **G.** GO Cellular Component (CC) analysis of DEGs. The top 5 terms were shown for both up and down-regulated genes. **H.** Enriched pathways (Metascape) of overlapping genes from human COPD patient ECs DEGs with DEGs from (D). **I.** GO BP, **J.** MF and **K**. CC analysis of common genes between human COPD patient lungs with DEGs in *Cry1/2* dKO ECs. Data are presented as mean ± SEM. ** p<0.1, *** p<0.001, and **** p<0.0001 *Cry1/2* si vs si_con (Two-way ANOVA followed by Bonferroni adjusted multiple comparisons). For **C**, only the asterisks for si-*Cry1/2* are shown.

To further explore the epithelial-specific effects of *Cry1/2* loss, we conducted RNA-sequencing on primary lung ECs isolated from WT and *Cry1/2* dKO mice at ZT4. Differential expression analysis revealed 501 upregulated and 369 downregulated genes in *Cry1/2* dKO ECs (Fig. 4D). Pathway enrichment analysis using Gene Ontology (GO) biological process (BP) analysis identified significantly upregulated pathways related to cilium movement and downregulated genes enriched in regulation of lamellipodium assembly and carbon dioxide transport (Fig. 4E). The molecular function (MF) analysis showed enrichment in tubulin binding and downregulation in genes involved in metal ion transporter activity (Fig. 4F). We also observed changes in cellular components (CC), including upregulation in structures such as the ciliary plasm and axoneme and downregulation in membrane microdomain and membrane raft (Fig. 4G). Furthermore, comparison of mouse DEGs with transcriptomic profiles from the lungs of human COPD patients^51^ revealed 123 overlapping genes (Fig. 4H). These genes were enriched in immune-related pathways including regulation of immune effector process, leukocyte migration and phagocytosis (Fig. 4I); in MF associated with peptide binding and immune receptor activity (Fig. 4J)j, and in structural components such as membrane microdomain, membrane raft, and phagocytic vesicles (Fig. 4K), suggesting that *Cry1/2*-deficient epithelial cells may acquire an aberrant proinflammatory phenotype that mirrors the inflammatory features observed in COPD. Together, these findings highlight a critical role for *Cry1/2* in maintaining alveolar epithelial homeostasis by regulating inflammatory response and cell proliferation, two processes central to COPD pathogenesis.

### *Cry1/2* depletion aggravates LPS induced inflammation in macrophages

In addition to ECs, inflammatory myeloid cells play an essential role in chronic inflammation in COPD ^52, 53^. To determine the role of *Cry1/2* in inflammatory cells, we isolated bone marrow derived macrophages (BMMs) from WT and *Cry1/2* dKO mice and measured the cytokine levels in BMMs either not treated or treated with 1 μg/ml LPS for 24 hours. Although the cytokine levels were not different between WT and *Cry1/2* dKO BMMs at baseline levels (Supplementary Fig. 4), LPS induced higher levels of *IL6*, matrix metalloproteinases 7 (*Mmp7),* and *Tnfa* in *Cry1/2* dKO BMMs compared to WT BMMs (Fig. 5A), suggesting that macrophage *Cry* deletion could contribute to lung inflammation. We next sought to investigate whether bone marrow (BM) cells with specific *Cry1/2* deletion aggravate LPS-driven COPD-like phenotype. For this purpose, BM cells were isolated from WT or *Cry1/2* dKO mice (CD45.2 positive strain) and transplanted into total-body irradiated (TBI) recipient mice (CD45.1 positive strain) (Fig. 5B). Eight Weeks after transplantation, flow cytometry was performed to examine the white blood cell types in the recipient mice. More than 95% of cells in the recipient mice were CD45.2, suggesting that donor mouse origin cells are the dominant population (Fig. 5C). Twelve weeks after bone marrow transplantation, BAL cells were collected from WT and *Cry1/2* dKO mice at ZT4–6 for transcriptomic profiling. RNA sequencing identified 414 upregulated and 280 downregulated genes in *Cry1/2* dKO BAL cells compared to WT controls (Fig. 5D). GO biological process analysis revealed that the upregulated genes were enriched in proinflammatory pathways, including positive regulation of response to virus, positive regulation of innate immune response, and response to biotic stimulus (Fig. 5E). In contrast, the downregulated genes were predominantly associated with cell cycle regulation, chromosome segregation, and nuclear division, suggesting impaired proliferative capacity (Fig. 5E). These findings indicate that *Cry1/2* deletion in hematopoietic cells promotes a heightened inflammatory state and may dysregulate immune cell turnover in the lung. Supporting this, cell-type enrichment analysis using xCell2 revealed that *Cry1/2* dKO BAL samples exhibited more prominent signatures for multiple immune cell types, including B cells, macrophages, monocytes, and neutrophils (Fig. 5F), further highlighting the proinflammatory and immune-activated phenotype resulting from *Cry1/2* loss in the bone marrow.

**Figure 5.**
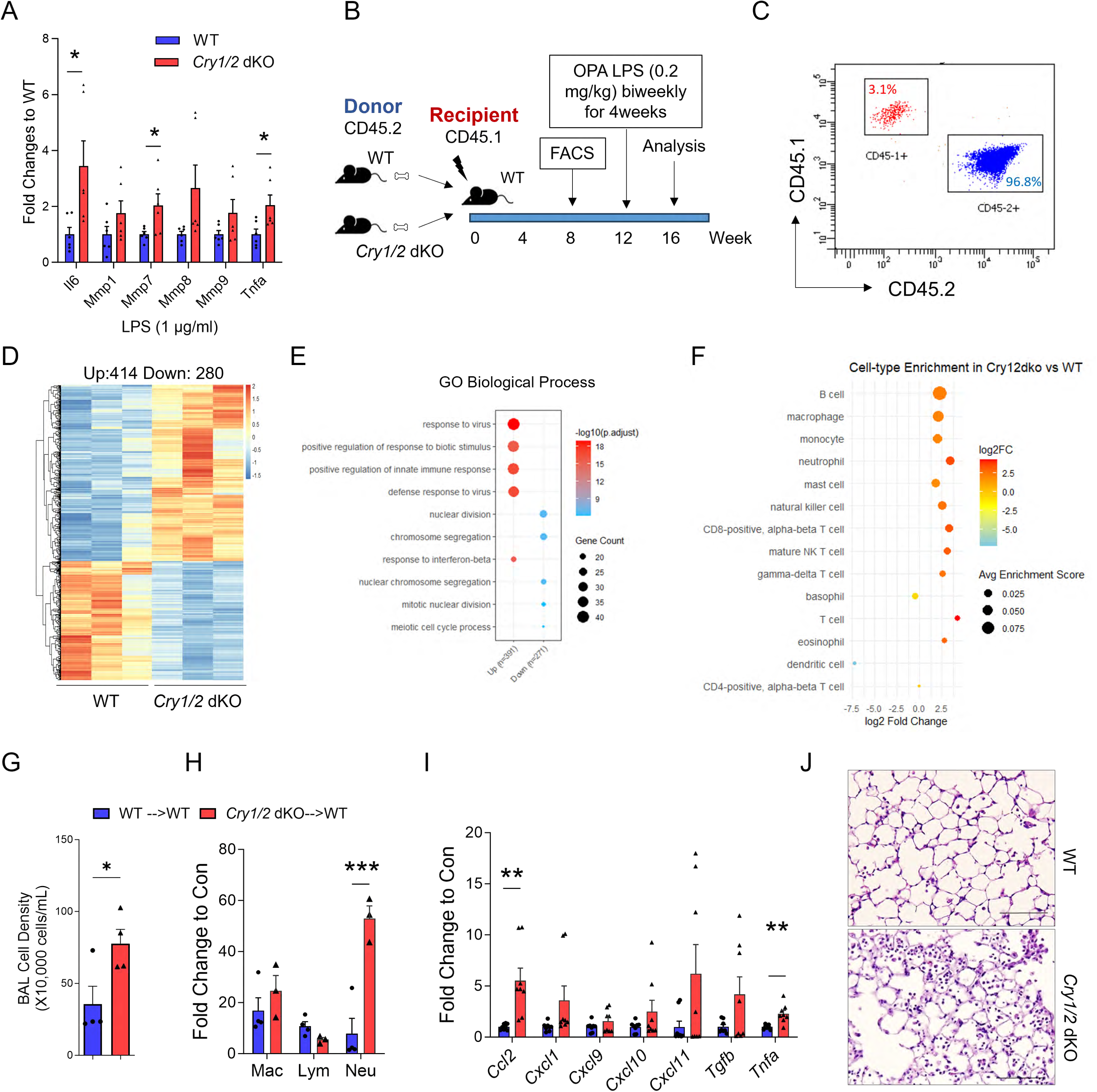
The role of *Cry1/2* in bone marrow (BM) cells. Bone marrow cells were isolated from 10-12 week old sex-matched WT or *Cry1/2* dKO mice and differentiated into macrophages using L929 conditioned medium. **A.** The cytokine mRNA levels were determined in bone marrow-derived macrophages (BMMs) from WT or *Cry1/2* dKO mice treated with 1 μg/ml LPS for 24 hrs. **B.** Experimental schematic: 3.5 × 10^6^ bone marrow (BM) cells from *Cry1/2* dKO and age- and sex-matched WT mice were retro-orbitally injected into lethally irradiated B6 CD45.1 mice on day 1. **C.** Eight weeks after transplantation, flow cytometry of blood cells was performed to determine the proportion of CD45.1+ and CD45.2+ cells. (**D-E**) Twelve weeks after transplantation, the BAL cells were collected from WT or *Cry1/2* dKO mice for RNA-sequencing. D. Heatmap showing 414 upregulated and 280 downregulated genes in the BAL cells from *Cry1/2* dKO mice. E. GO analysis identified the top 5 upregulated or downregulated biological processes in *Cry1/2* dKO mice. **F.** XCell2 identified enriched cells from the DEGs of *Cry1/2* dKO mice. (**G-J**) Twelve weeks after transplantation, mice were treated with LPS (0.2 mg/kg) biweekly for 4 weeks and analyzed on day 28. **G.** Total BAL cell count. **H.** The number of macrophages, lymphocytes, and neutrophils in BAL. **I.** Cytokine transcript levels in the whole lungs. **J.** Representative images of H&E-stained slides (scale bar=100 μm). Data are presented as mean ± SEM. * p<0.05, ** p<0.01, *** p<0.001 based on Bonferroni adjusted t-test.

Next, we treated the recipient mice with chronic LPS (0.2 mg/kg body weight twice a week) starting at 12 weeks after BM transplantation. Mice transplanted with *Cry1/2* dKO BM showed significantly increased total BAL cells (Fig. 5G), specifically neutrophils (Fig. 5H), elevated *Ccl2* and *Tnfa* levels (Fig. 5I), and severe airway inflammation compared to WT BM transferred mice (Fig. 5J). These findings indicate that CRY1/2 proteins in BM-derived cells play a protective role against LPS-induced lung inflammation. However, no further airspace enlargement was observed in mice receiving *Cry1/2* dKO BM compared to those receiving WT BM (Fig. 5J), suggesting that *Cry1/2* deletion in BM alone may not be sufficient to drive emphysema pathology. Thus, *Cry1/2* expression in both epithelial and immune compartments may be required to maintain lung homeostasis and prevent emphysematous remodeling.

### Nobiletin (NOB) attenuates the emphysema-like phenotype and inflammation in *Cry1/2* dKO mice

We previously identified Nobiletin as a clock-enhancing molecule (CEM) from high-throughput chemical screening, acting to directly interact and activate the core clock component ROR nuclear receptors ^54^. Our functional studies demonstrated significant *in vivo* activity and disease relevance of NOB ^55, 56, 57, 58, 59^. To determine whether NOB could rescue the lung pathology observed in *Cry1/2* dKO mice, we initiated a NOB-containing diet at weaning (postnatal day 21) and continued treatment until the mice reached 20 weeks of age (Fig. 6A). NOB treatment significantly reduced airspace enlargement in *Cry1/2* dKO mice, as evidenced by histological analysis and mean linear intercept measurements (Fig. 6B, 6C). To further explore the molecular effects of NOB on alveolar epithelial cells (ECs), we isolated primary alveolar ECs from WT mice and *Cry1/2* dKO mice fed either a regular diet (RD) or a NOB diet at ZT4 and performed RNA-sequencing to assess transcriptional changes. We identified 501 upregulated and 369 downregulated genes in *Cry1/2* dKO ECs compared to WT controls. Interestingly, 390 upregulated and 81 downregulated genes were restored to WT levels following NOB treatment (Fig. 6D). The GO biological processes analysis suggested that rescued genes were strongly enriched in processes related to cilium movement, cilium organization, and microtubule-based motility (Fig. 6E). The molecular function analysis showed enrichment in microtubule binding, metal ion transporter activity, and chemoattractant receptor activity (Fig. 6F). In terms of cellular components, there was a strong enrichment in structures such as the axoneme, motile cilium, and plasma membrane-bounded cell projection cytoplasm (Fig. 6G). These findings are consistent with the biological process results and indicate that NOB promotes recovery of apical epithelial structures required for coordinated ciliary beating and surface integrity. Upregulation of genes involved in cilium movement was further confirmed by RT-qPCR, and these genes were fully rescued by NOB treatment (Fig. 6H). Additional rescued genes include complement components (*C2, C4b*), chemokines (*Cxcl12, Cxcl14, Cxcl17*), interleukins (*IL16*), and MMPs (*Mmp10, Mmp2, Mmp3*) (Fig. 6I). NOB also attenuated the downregulation of surfactant proteins (*Sftpb* and *Sftpc*) observed in *Cry1/2* dKO alveolar ECs (Fig. 6I), suggesting that it may either mitigate alveolar EC death or improve the alveolar EC proliferation.

**Figure 6.**
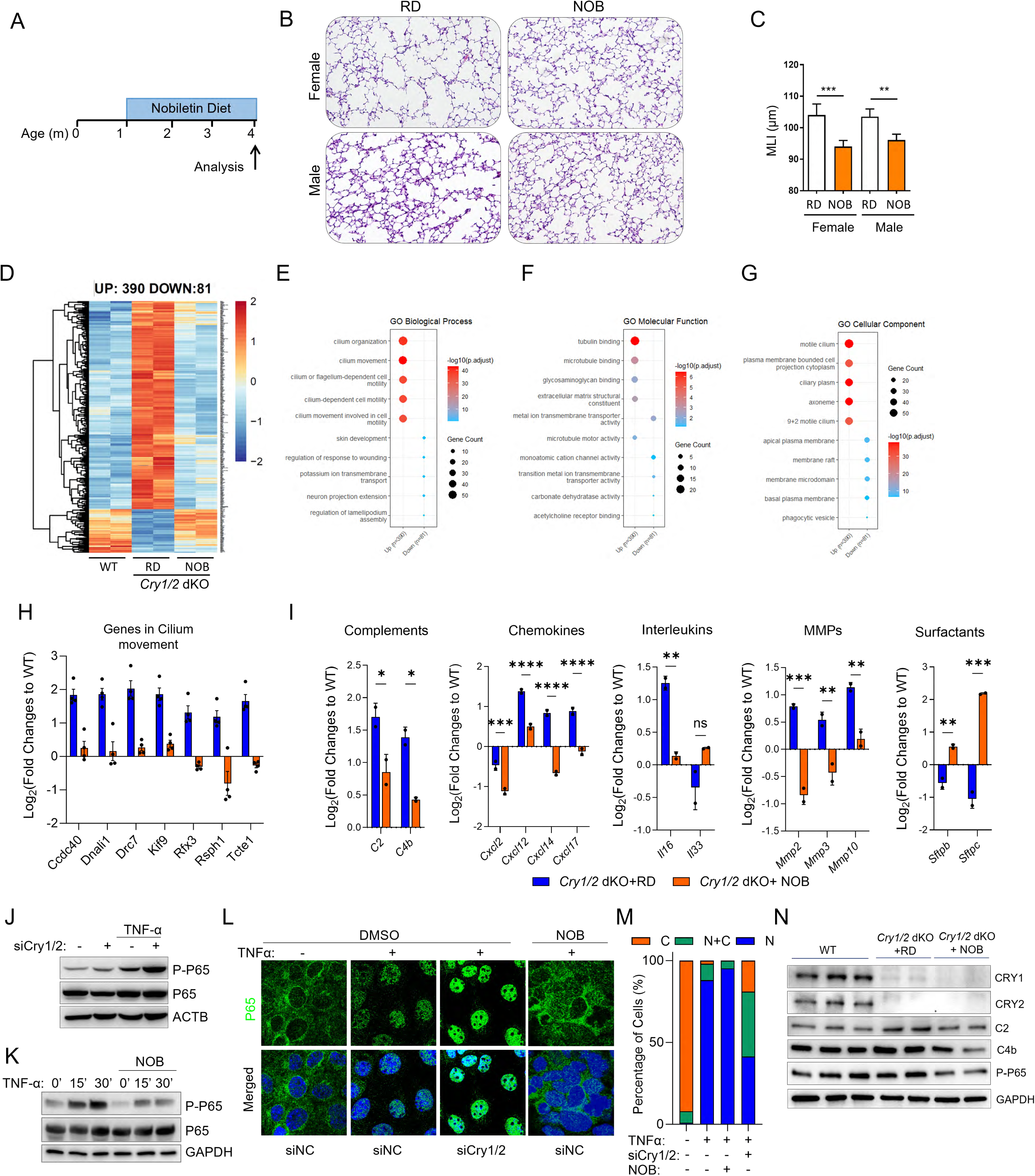
NOB attenuates emphysema and inflammation in *Cry1/2* dKO mice. **A.** Experimental schematic: *Cry1/2* dKO mice were fed with regular (RD) or 1% NOB-containing diets starting at 1 month old and analyzed at 4 months. **B.** Representative images of H&E-stained lung sections collected from mice fed with a RD or NOB diet. **C.** MLI quantification of H&E-stained lung sections from male or female *Cry1/2* dKO fed with RD or NOB diets. **D.** RNA-seq was performed using primary lung ECs isolated from the WT mice, or *Cry1/2* dKO mice fed with RD or NOB diet. Heatmap of RNA-seq data of lung ECs from different treatment groups. **E.** GO BP analysis of DEGs that were rescued by NOB treatment in *Cry1/2* dKO lung ECs. **F.** GO MF analysis of rescued DEGs. **G.** GO CC analysis of rescued DEGs. **H.** RT qPCR confirmed the upregulation and rescue of genes involved in cilium movement. **I.** The transcript expression changes of important differentially expressed chemokines, cytokines, *Mmps*, and surfactant proteins from RNA-seq data. Data were presented as fold changes to WT. **J.** MlE12 cells were transfected with control siRNA or siRNA targeting *Cry1* and *Cry2*. 36 h after transfection, cells were treated with 5 ng/ml TNF-α for 15 min. Phosphorylated P-65 and P65 levels were determined with Western blots. **K.** MLE12 cells were treated with 20 µM NOB for 24 h, followed by 5 ng/ml TNF-α for 0, 15, or 30 mins. Phosphorylated P-65 and P65 levels were determined with Western blots. **L.** MLE12 cells were transfected with control siRNA, or siRNA targeting *Cry1* and *Cry2*, and treated with 20 µM NOB for 24 h before TNF-α stimulation (15 mins). Representative confocal images of p65 (green) are shown (×400 magnification; scale bar = 10 µm). **M.** The percentage of cells with p65 localized in the cytoplasm (C), both cytoplasm and nucleus (C+N), or nucleus (N) was quantified for each treatment condition. **N.** Western blot of *CRY1/2* and COPD-associated proteins in primary lung ECs from different treatment groups. Data are presented as mean ± SEM. P-value was calculated from a two-tailed t-test for H and I, * p<0.05, ** p<0.01, *** p<0.001, and **** p<0.0001 in RD vs in *Cry1/2* dKO samples.

Next, we investigated the molecular mechanisms underlying the effects of NOB in lung ECs. Previous studies have shown that CRY proteins reduce cAMP production, leading to downstream suppression of protein kinase A (PKA) and NF-κB signaling ^60^. Importantly, our recent work demonstrated that NOB activates RORs and their target gene *IkBα*, resulting in the suppression of the NF-κB signaling pathway in triple-negative breast cancer ^58^. To investigate the impact of *Cry1/2* on the NF-κB pathway, MLE12 cells were transfected with control siRNA or siRNA targeting *Cry1* and *Cry2*, followed by treatment with TNF-α for 15 min. Western blot analysis showed that *Cry1/2* knockdown has a minimal effect on p65 phosphorylation in cells without TNF-α; however, upon TNF-α stimulation, *Cry1/2* knockdown highly increased phospho-p65 levels (P-P65), the activated form of the NF-κB subunit p65, which indicates NF-κB pathway activation (Fig. 6J). Next, to examine the mechanistic role of NOB in NF-κB regulation, MLE12 cells were pretreated with 20 µM NOB for 24 h, followed by TNF-α stimulation. Western blot analysis demonstrated that TNF-α activated NF-κB signaling by inducing phosphorylation of p65 at as early as 15 min, with the effect sustained at 30 min, whereas NOB significantly attenuated phospho-p65 levels at both time points (Fig. 6K). We further examined the effects of NOB and *Cry1/2* depletion on p65 nuclear localization by immunofluorescent staining. TNF-α treatment robustly induced p65 nuclear translocation, which was further enhanced by knockdown of *Cry1/2* and effectively inhibited by NOB pretreatment (Fig. 6L and 6M). Consistent with these findings, we confirmed that *Cry1/2* deletion modestly increased phospho-p65 protein levels in primary isolated *Cry1/2* dKO alveolar epithelial cells (Fig. 6N). Importantly, NOB markedly reduced phospho-p65 levels in *Cry1/2* dKO epithelial cells and attenuated the upregulation of proinflammatory proteins induced by *Cry1/2* deficiency, including C2 and C4b (Fig. 6N). Overall, these findings highlight a potential role of circadian modifiers such as NOB in protecting lungs from COPD-related injury by mitigating NF-κB activity and limiting proinflammatory signaling in epithelial cells.

## Discussion

COPD is a complex and progressive respiratory disorder characterized by persistent inflammation, alveolar destruction, and impaired tissue repair. Our study provides *in vivo* and *in vitro* evidence that CRYs are critical circadian repressors that maintain lung integrity by regulating epithelial cell proliferation and inflammatory response. The loss of *Cry1/2* sensitizes the lung to spontaneous emphysema-like pathology, while the pharmacologic clock modifier NOB can attenuate these detrimental effects. These findings illustrate the molecular interplay between the circadian clock and lung disease and support the emerging concept of targeting circadian pathways as a novel strategy for treating COPD and other chronic inflammatory conditions.

Although prior studies have implicated circadian rhythms in the regulation of lung physiology, the underlying molecular mechanisms linking circadian disruption to COPD pathogenesis remain poorly defined. Our study pinpoints the core clock components CRY1 and CRY2 as key regulators of pulmonary homeostasis. In *Cry1/2* dKO mice, we observed spontaneous development of emphysema-like pathology, including alveolar enlargement, increased lung compliance, decreased elasticity and heightened inflammatory cell infiltration in the absence of external insults such as CS or pollutants. These findings highlight a more profound and intrinsic role for *Cry1/2* in lung homeostasis, in contrast to previous studies where other circadian mutant models, such as *Bmal1*-deficient mice, only demonstrated exaggerated inflammation and emphysematous changes following exposure to chronic CS in combination with influenza A virus infection ^38^. *Rev-erbα* KO mice showed heightened inflammatory responses accompanied by an increase in cellular senescence markers in response to LPS ^61^; however, lung functional assays were not employed to assess disease progression. In comparison, our study provides direct physiological evidence using flexiVent-based lung function measurements that *Cry1/2* dKO mice exhibit significantly increased lung compliance and reduced elasticity. This early-onset phenotype underscores the essential role of *Cry1/2* in protecting the lung from spontaneous degenerative changes.

Consistent with this notion, we also found that CRY2 protein levels were significantly reduced in lungs from mice exposed to chronic LPS, a model that mimics chronic inflammation observed in COPD. This observation is consistent with reports showing reduced expression of core clock genes, including CRY1 and CRY2 in lung tissues from COPD patients ^62, 63^. RNA-seq data from bronchial brushings of COPD patients revealed that higher expression of core clock genes (*Per1-3*, *Cry1/2* ), based on a composite CCG score, was associated with lower expression of proinflammatory genes, including *CXCL8*, *IL1A*, *IL1B*, and *TGFB1* ^64^. Similar alterations have been documented in various animal models, including those exposed to CS, where expression of *Cry1/2, Pers, Bmal1,* and *Clock* is dysregulated alongside reduced locomotor activity ^37, 65, 66^. Additionally, chronic CS exposure together with influenza A virus infection exacerbates lung inflammation and promotes emphysema in mice in association with impaired pulmonary rhythm and disrupted normal timing of circadian gene expression, including *Cry1*, *Per1*, *Clock*, *Baml1*, and *Rev-erb-α* ^38^. These data are further supported by the evidence that inflammation itself can modulate clock gene expression. For instance, LPS injection leads to suppression of *CLOCK, CRY1/2, PER1-3, CSNK1ε, RORα, and REV-ERBα* expression in human peripheral blood lymphocytes, neutrophils, and monocytes ^67^. Moreover, various cytokines, i.e., IFN-γ, TNF-α, and IFN-α, can alter clock gene expression ^68^. Collectively, these findings reinforce the relationship between circadian disruption and COPD.

Our study further demonstrates that *Cry1/2* deficiency amplifies inflammatory signaling within the lung epithelium and macrophages. Neutrophil counts were significantly elevated in the BAL of *Cry1/2* dKO mice compared to WT controls, which may partially contribute to the emphysema-like phenotype ^69^. Knockdown of *Cry1/2* in alveolar epithelial cells led to upregulation of *Ccl2*, *Cxcl1*, *Tnfa*, and *Il8* (*Cxcl15* in mice), as well as impaired cell expansion. Transcriptomic analysis of primary lung epithelial cells from *Cry1/2* dKO mice revealed altered expression of genes involved in immune activation, overlapping with gene signatures from COPD patient epithelium. In addition to epithelial cells, our studies also demonstrated that macrophages lacking *Cry1/2* produce higher levels of pro-inflammatory cytokines in response to LPS stimulation. Our results are consistent with previous findings showing that bone marrow-derived macrophages and fibroblasts from *Cry1/2* dKO mice exhibit elevated expression of proinflammatory cytokines (IL-6, TNF-α, iNOS, CXCL1), and that CRY-deficient animals have heightened inflammatory responses following LPS exposure ^60^. Mechanistically, CRY proteins have been shown to bind and inhibit adenylyl cyclase activity, thereby reducing cAMP production and downstream suppression of protein kinase A (PKA) and NF-κB signaling. CRY1/2 can also attenuate TNF-α–mediated inflammation in fibroblast-like synoviocytes by dampening both PKA and NF-κB pathways ^70^. *Cry1/2* double knockout mice display autoimmune-like features, including elevated B2 B cell populations in the peritoneal cavity and increased leukocyte infiltration in the lungs and kidneys^71^. Conversely, overexpression of CRY1 has been shown to suppress sleep deprivation–induced vascular inflammation by inhibiting cAMP/PKA signaling and NF-κB activation ^72^. Our findings support and expand upon this concept by showing that *Cry1* and *Cry2* exert anti-inflammatory functions not only in immune cells but also in lung epithelial cells, two central players in COPD progression.

In addition to their anti-inflammatory roles, *Cry1/2* are also implicated in regulating epithelial proliferation. In our study, silencing *Cry1* or *Cry2* in lung epithelial cells led to significantly suppressed cell proliferation, suggesting a role in maintaining epithelial integrity. Supporting this, studies have shown that disruption of *Bmal1 or Cry1/2* severely impairs lung epithelial regeneration, as tracheal organoids derived from *Bmal1* KO or *Cry1/2* dKO mice showed drastically reduced growth, highlighting a critical role for circadian proteins in maintaining lung repair capacity ^73^. Similarly, CRY1 depletion is reported to induce ovarian granulosa cell senescence ^74^. CRY2 can interact with mRNA regulators (e.g., Bclaf1 and cyclin D1), and its loss triggers premature cell cycle exit, disrupting muscle regeneration ^75^. Given the strong impact of alveolar epithelial cell death/senescence on the pathogenesis of emphysema, our findings raise the possibility that *Cry1/2* deficiency compromises epithelial renewal, thereby increasing susceptibility to tissue degeneration and airspace enlargement. RNA-seq analyses of primary isolated lung alveolar epithelial cells revealed that the enriched gene signatures in *Cry1/2* dKO epithelial cells include cilium organization, cilium movement, and microtubule-based motility. These biological processes are typically associated with airway multiciliated cells rather than alveolar epithelial cells. However, given that *Cry1/2* knockdown alveolar epithelial cells show attenuated proliferation and *Cry1/2* dKO mice display clear signs of alveolar airspace enlargement, the upregulation of ciliogenesis-related programs may represent a stress-induced aberrant epithelial regeneration ^76, 77^. Alveolar epithelial cell death or senescence in the *Cry1/2* dKO lung may trigger compensatory recruitment or differentiation of basal-like progenitor cells from adjacent bronchiolar regions ^76, 78^. These progenitors, in response to injury signals and loss of alveolar niche integrity, may undergo a transitional or intermediate epithelial state, representing a phenomenon increasingly recognized in lung injury and repair models. These transitional cells may transiently express airway-associated genes, including those involved in ciliogenesis, as part of an aberrant or incomplete repair process.

Importantly, our study highlights the therapeutic potential of targeting circadian pathways using NOB, a natural flavonoid previously identified as a clock-modulating molecule ^54^. Our studies showed that NOB administration significantly mitigated lung inflammation and alveolar destruction in *Cry1/2* dKO mice. Transcriptomic analyses revealed that NOB reversed many of the gene expression changes observed in CRY-deficient alveolar epithelial cells, including restoration of surfactant protein expression and suppression of cytokines, chemokines, and matrix metalloproteinases. These data are consistent with recent findings showing that NOB represses inflammatory pathways in other tissues, including liver, muscle, and vasculature ^55, 56, 79, 80, 81^. Mechanistically, our previous findings suggest that NOB may exert its protective effects in the lung through ROR-mediated transcriptional activation of IκBα, an endogenous inhibitor of NF-κB, thereby suppressing NF-κB–driven inflammation even in the absence of CRY1/2 ^58^. CRY1/2 are known to inhibit adenylyl cyclase activity, resulting in lower intracellular cAMP levels, reduced PKA activation, and diminished phosphorylation of NF-κB subunit p65 ^60^. In *Cry1/2*-deficient mice, loss of this inhibition led to elevated cAMP-PKA signaling and enhanced p65 phosphorylation, which promotes the transcription of proinflammatory genes. Notably, NOB treatment of *Cry1/2* dKO mice reduced p65 phosphorylation, suggesting that NOB can re-establish anti-inflammatory signaling after CRY1/2 depletion. These data collectively position NOB and other circadian interventions as promising therapeutic strategies for inflammatory lung diseases such as COPD.

In summary, our study revealed an important role of *Cry1/2* in maintaining lung integrity by regulating epithelial cell proliferation and inflammatory response. Moving forward, further studies will aim to dissect the mechanisms by which CRY1/2 regulate epithelial cell proliferation and inflammatory response. We will also investigate the cell-specific functions of CRY1/2 in various pulmonary compartments, such as alveolar epithelial cells, macrophages, and neutrophils, to better define their contributions to COPD pathogenesis. Additionally, testing the impact of *Cry1/2* deficiency in other established models of COPD, such as CS exposure and porcine pancreatic elastase (PPE)-induced emphysema, will provide valuable insights into the generalizability and robustness of CRY-mediated protection. Finally, the development and evaluation of other circadian-targeting therapies, such as synthetic ROR agonists and REV-ERB modulators, may pave the way for precision-based interventions that harness the circadian clock to restore lung homeostasis and mitigate COPD progression.

## ACKNOWLEDGMENTS

We would like to thank Dr. Farrah Kheradmand at Baylor College of Medicine for her thoughtful advice on this project, and Kaori Ono for her assistance with mouse maintenance.

## AUTHOR CONTRIBUTIONS

S.Y. and T.M. designed the study. T.M., S.Y., C.H., J.L., J.W., H.L., Y.W., and K.Y. performed the described experiments. N.K. and S.K. conducted bioinformatics analyses. T.M. and S.Y. drafted the manuscript. J.L.A. and Z.C. reviewed and revised the manuscript.

