## Supplemental materials for "Cryptochrome Loss Drives COPD-like Lung Pathology through Disrupted Alveolar Epithelial Proliferation and Immune Homeostasis"

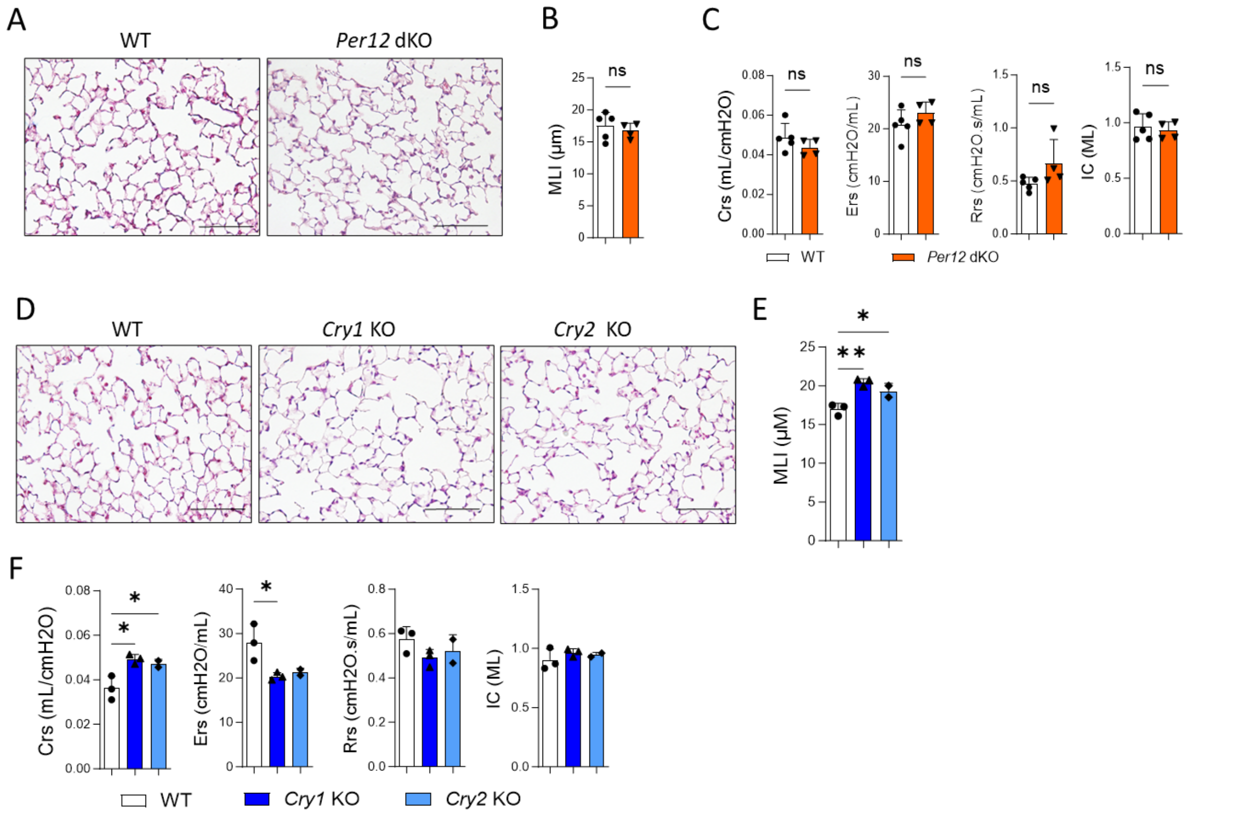


**Supplementary Figure 1: Lung sections of *Per1/2* dKO and *Cry1* and *Cry2* single KO mice**. Mouse lung sections were collected from 12-18-week-old mice. (**A** and **D**) Representative images of hematoxylin and eosin (H&E) stained (**A**) Female WT and *Per1/2* dKO mice and (**D**) Male WT, *Cry1* KO or *Cry2* KO mice. (**B** and **E**) Quantification of airspace enlargement by mean linear intercept (MLI) analysis in (**B**) WT and *Per1/2* dKO mice and (**E**) WT, *Cry1* KO, or *Cry2* KO mice. (C and F) Lung function measurements obtained using the flexiVent system, including compliance, elasticity, and inspiratory capacity in (**C**) WT and *Per1/2* dKO mice and (**E**) WT, *Cry1* KO or *Cry2* KO mice. Scale bar=200 μM. P-value was calculated from a two-tailed t-test to compare between the WT and *Per1/2* dKO mice, and it was calculated from one-way ANOVA followed by Dunnett comparison among WT, *Cry1* KO or *Cry2* KO mice. * p<0.05 and ** p<0.01.

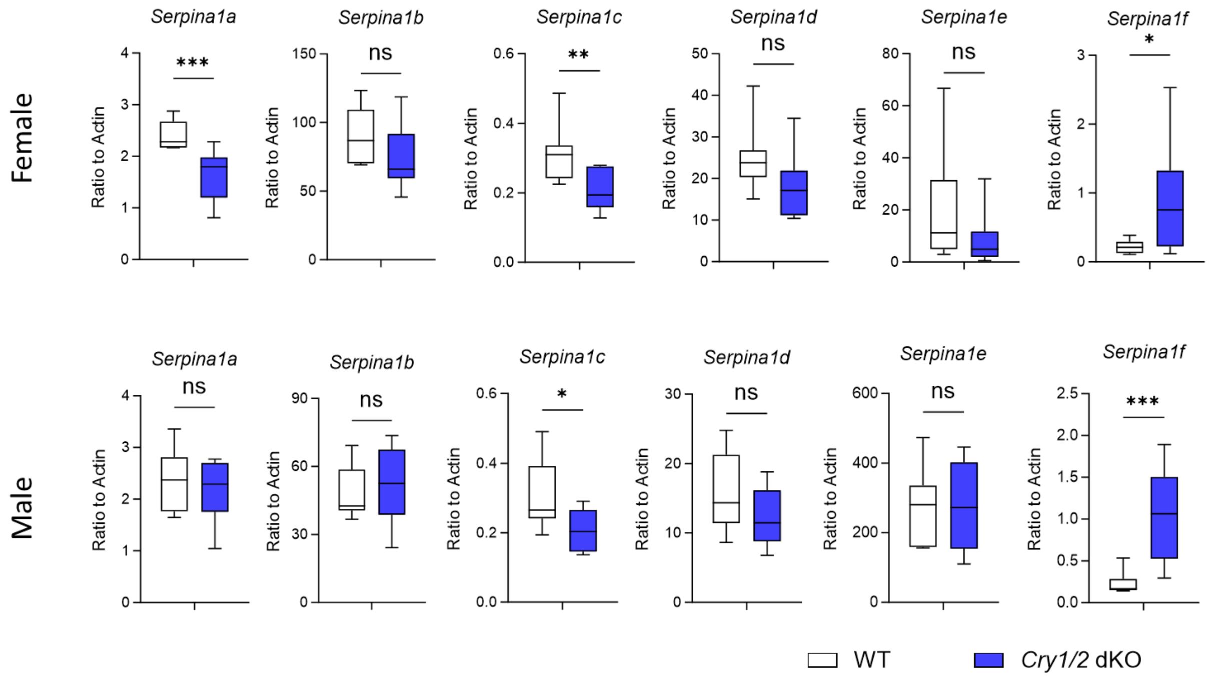


**Supplementary Figure 2: The transcript levels of A1AT genes in *Cry1/2* dKO mouse livers**. Mouse livers were collected from 10-12-week-old male or female WT and *Cry1/2* dKO mice at ZT12. RT qPCR was performed to determine the levels of Alpha-1 antitrypsin (A1AT) encoding Serpina1 genes, including *Serpina1a*, *Serpina1b*, *Serpina1c*, *Serpina1d*, *Serpina1e*, *Serpina1f*, and other serpin genes belonging to the serine protease inhibitor superfamily, including *Serpina3g*, *Serpina3k*, *Serpina3n*, *Serpinab1a*, *Serpind1,* and *Serping1*. P-value was calculated from a two-tailed t-test, * p<0.05, ** p<0.01, and *** p<0.001 in RD vs in *Cry1/2* dKO.

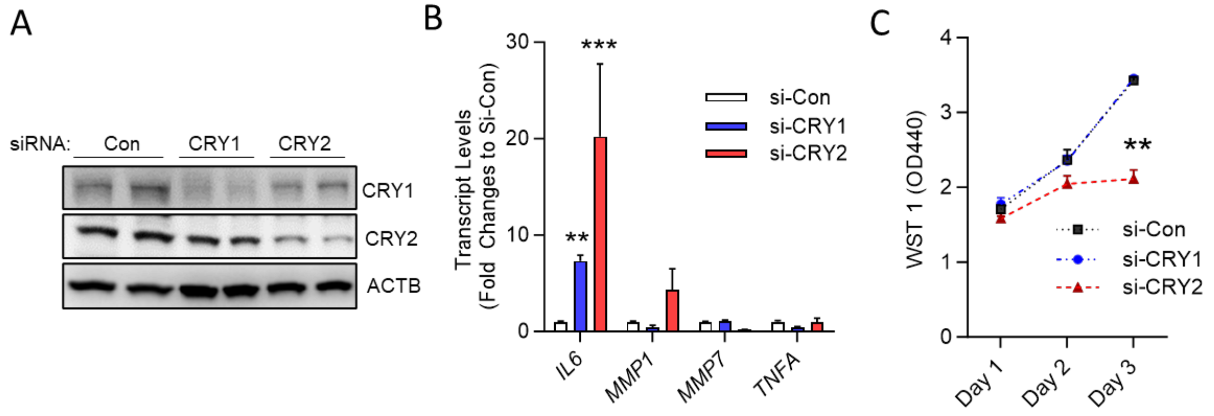


**Supplementary Figure 3: The impact of *CRY1/2* knockdown in A549 cells.** Human lung epithelial A549 cells were transfected with 50 pmole/ml control, CRY1, or CRY2 siRNA on day 0 and day 1. **A.** Western blot showed decreased CRY1/CRY2 protein levels on 2 days post-transfection. **B.** The mRNA levels of cytokines, including Interleukin-6 (*IL6*), matrix metalloproteinase 1 and 7 (MMP1 and MMP7), and *TNFA,* were determined with RT-qPCR. C. Cell proliferation was measured with WST1 assay 1-, 2-, or 3-days post siRNA transfection. n=4 ** p<0.1 and *** p<0.001, si_*Cry1/2* vs si_con (Two-way ANOVA followed by Bonferroni adjusted multiple comparisons).

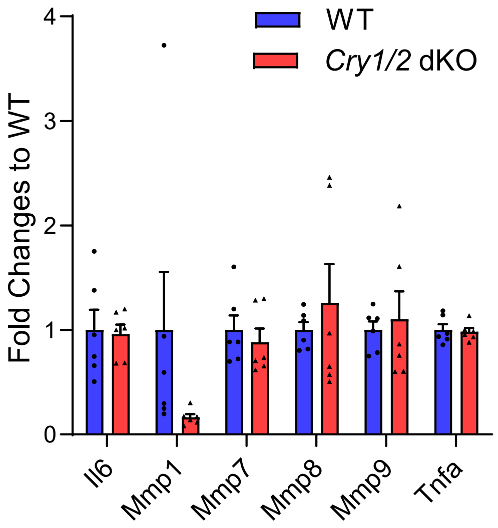


**Supplementary Figure 4: The baseline cytokine levels in bone marrow-derived macrophages (BMMs) from WT and *Cry1/2* dKO mice.** Bone marrow cells were isolated from 10-12 week old sex-matched WT or *Cry1/2 dKO* mice and differentiated into macrophages using L929 conditioned medium. The transcript levels of cytokines were determined in BMMs from WT or *Cry1/2 dKO* mice without any treatment.

**Supplementary Table 1: RT qPCR primers**

| Gene | Forward primer | Reverse Primer |
|---|---|---|
| Homo_ACTA1 | CATGTACGTTGCTATCCAGGC | CTCCTTAATGTCACGCACGAT |
| Homo_CXCL1 | ACTGCTGCTCCTGCTCCT | CGATGATTTTCTTAACTATGGG |
| Homo_IL6 | AATTCGGTACATCCTCGACGG | TTGGAAGGTTCAGGTTGTTTTCT |
| Homo_IL8 | CTTTCCACCCCAAATTTATCAAAG | CAGACAGAGCTCTCTTCCATCAGA |
| Homo_MMP1 | AAAATTACACGCCAGATTTGCC | GGTGTGACATTACTCCAGAGTTG |
| Homo_MMP7 | GAGTGAGCTACAGTGGGAACA | CTATGACGCGGGAGTTTAACAT |
| Homo_TNFA | GAGGCCAAGCCCTGGTATG | CGGGCCGATTGATCTCAGC |
| Mus_Acta1 | GGCTGTATTCCCCTCCATCG | CCAGTTGGTAACAATGCCATGT |
| Mus_Ccl2 | TTAAAAACCTGGATCGGAACCAA | GCATTAGCTTCAGATTTACGGGT |
| Mus_Cxcl1 | CTGCACCCAAACCGAAGTC | AGCTTCAGGGTCAAGGCAAG |
| Mus_Cxcl10 | CCAAGTGCTGCCGTCATTTTC | GGCTCGCAGGGATGATTTCAA |
| Mus_Cxcl11 | GGCTTCCTTATGTTCAAACAGGG | GCCGTTACTCGGGTAAATTACA |
| Mus_Cxcl9 | TCCTTTTGGGCATCATCTTCC | TTTGTAGTGGATCGTGCCTCG |
| Mus_E-cad | CAGGTCTCCTCATGGCTTTGC | CTTCCGAAAAGAAGGCTGTCC |
| Mus_Homo_18S | GTAACCCGTTGAACCCCATT | CCATCCAATCGGTAGTAGCG |
| Mus_IIL8(Cxcl15) | TCGAGACCATTTACTGCAACAG | CATTGCCGGTGGAAATTCCTT |
| Mus_Mmp1/13 | CTTCTTCTTGTTGAGCTGGACTC | CTGTGGAGGTCACTGTAGACT |
| Mus_Mmp10 | GAGCCACTAGCCATCCTGG | CTGAGCAAGATCCATGCTTGG |
| Mus_Mmp7 | CTGCCACTGTCCCAGGAAG | GGGAGAGTTTTCCAGTCATGG |
| Mus_Mmp9 | GAGACGGGTATCCCTTCGAC | TGACATGGGGCACCATTTGAG |
| Mus_Serpina1a | TATCTCCTCCAGCCATGCAA | GAGTCATTGAGCATGTCCTCTT |
| Mus_Serpina1b | CAGCAGCTACAGTCTTTGAAGC | GGGTGGTCATTTATGTGTGGG |
| Mus_Serpina1c | TCCATCTTGGCTCCCACTAC | CCTGAACATCCTCAGCCAGA |
| Mus_Serpina1d | GATGGATTACGCAGGCAACA | TGGGGATATGGATCTGAGCAT |
| Mus_Serpina1e | TATGCCCCCTATCTTGCACTT | GCCCGTGTTTAATGGAAAGGA |
| Mus_Serpina1f | ATGACAACACCCTTTTCCTCCC | GCAGCAATGACTCTAATCGGG |
| Mus_Serpina3g | GGCCTGAAAGAGAGCACATT | CATTCGGGTCAAAGGGGTTC |
| Mus_Serpina3k | TGAGGAGCTATCGTGCTCTGT | GCCTGTAGTTACTAGCGATGGA |
| Mus_Serpina3n | ATTTGTCCCAATGTCTCCCAA | TGCCTATCTTGGCTATAAAGGGG |
| Mus_Serpinb1a | ACATCCATTCACGCTTCCAAA | GCCCAAGTCAGCACCATACAT |
| Mus_Serpind1 | GATGCCGTTTCCCCGACAG | TCAGGACTCGGTAAAGATTCAA |
| Mus_Serping1 | TAGAGCCTTOTCAGATCCCGA | ACTCGTTGGCTACTTTACCCA |
| Mus_Sftpd | TGAAGAGCCTCTCGCAGAGATC | TAGGACCTGGTTTGCCTTGAGG |
| Mus_Tgfb1 | CTCCCGTGGCTTCTAGTGC | GCCTTAGTTTGGACAGGATCTG |
| Mus_Tnfa | CACCACCATCAAGGACTCAA | TCCAGCCTCATTCTGAGACA |
